# fourSynergy: Ensemble-based interaction calling on 4C-seq data using gradient-free optimization

**DOI:** 10.64898/2026.05.27.728108

**Authors:** Sophie-Marie Wind, Lucas Plagwitz, Jonas Dix, Gero Heidtmann, Dominik Heider, Carolin Walter

## Abstract

**Motivation:** Chromatin organization plays a crucial role in gene regulation and is associated with various severe diseases like cancer. Since chromatin changes are potentially reversible, a deeper understanding of the alterations needs to be harnessed for the development of new therapies. Circular Chromosome Conformation Capture Sequencing (4C-seq) is a sequencing technique enabling the identification of chromatin interactions between genes and regulatory elements. This work aims to develop an ensemble algorithm that utilizes synergies among available 4C-seq tools, which in turn allows to achieve superior predictive performance in interaction calling.

**Results:** We employed existing 4C-seq algorithms using a weighted-voting approach. By optimizing the tool weights according to various predictive metrics using gradient-free optimization strategies, we demonstrate the potential of combining multiple 4C-seq analysis tools for interaction calling. Our results indicate that a weighted-voting based ensemble approach can outperform individual algorithms in various datasets. Although the optimal solutions differ across the 4C-seq datasets, we successfully identified global solutions that outperform the individual algorithms for all datasets analyzed.

**Availability:** https://github.com/sophiewind/fourSynergy, https://github.com/sophiewind/fourSynergy_pip

**Supplementary information:** Supplementary data are available at *Journal Name* online.

## Introduction

The spatial organization of chromatin is crucial for gene regulation and is associated with various severe diseases, such as cancer. Recent studies have shown that aberrations in chromatin conformations contribute to expression changes in acute myeloid leukemia (Schuetzmann et al., 2018; Ghasemi et al., 2021). Furthermore, dexamethasone treatment for breast cancer was shown to affect the spatial organization of the genome (Hoffman et al., 2022). Deeper insights into the chromatin landscape hold tremendous potential in the field of cancer research, since alterations in chromatin architecture are potentially reversible and, therefore, can be harnessed to develop new tumor therapies.

The development of chromosome conformation capture (3C)-based methods revolutionized the understanding of the spatial structure of the genome, providing insights into the interplay of chromatin organization and transcriptional regulation (Krijger et al., 2020). Among these techniques, Circular Chromosome Conformation Capture Sequencing (4C-seq) stands out for its ability to focus on a specific locus of interest, known as the viewpoint or bait, and generate a high-resolution map of all interactions with the entire genome (“one vs all”) (Han et al., 2018). It achieves a higher resolution than Hi-C while reducing sequencing costs. This makes 4C-seq a crucial technique for analyzing the spatial organization of the genome and for elucidating the interplay between genes and regulatory elements (Krijger et al., 2020).

The analysis of 4C-seq data is highly complex due to several technical limitations. The signal density is typically highest around the viewpoint (“near-bait”) and decreases with its distance from the viewpoint (“far-cis”)(Walter et al., 2019). Additionally, there should be more intra-than interchromosomal interaction, leading to a high coverage on the viewpoint chromosome and comparable low coverage on other chromosomes (“trans”). This is in line with the chromosome territory model. Furthermore, PCR bias can lead to an overrepresentation of specific regions (Raviram et al., 2016). Identifying interactions while complying with the specific requirements of 4C-seq data is a challenging task for which a number of algorithms have been developed. r3Cseq is an R/Bioconductor package developed by Thongjuea et al. for analyzing 4C-seq experiments (Thongjuea et al., 2013). Potential interactions are detected using adjustable length windows measured in base pairs (bp). For each potential interaction, p-values are calculated by comparing the residuals. fourSig is a software suite written in R and Perl. It is particularly interesting for far-cis analysis because of its optional masking of viewpoints (Williams Jr et al., 2014). peakC is a non-parametric 4C-seq analysis tool written in R and based on rank products. If replicates are available, the incorporation of replicate information helps to reduce the false positive rate and thus to improve precision. The tool focuses primarily on the detection of near-cis interactions (Walter et al., 2019; Geeven et al., 2018). 4C-ker (Raviram et al., 2016) is a Hidden-Markov Model (HMM) based tool and supports both single sample and differential analysis. Since the differential analysis is based on DESeq2, it relies on replicates. The tools classifies the genomic region into high- and low-interacting regions as well as regions without interactions (Raviram et al., 2016). Another 4C-seq analysis package is FourCSeq. It was provided as an R/Bioconductor package and supports single-sample and differential analysis. It identifies peaks based on fitting curves to fragment read counts and calculating z-scores from residuals (Klein et al., 2015). However, the package is deprecated now.

A benchmarking of the existing 4C-seq tools reveals that none of the tools performs adequately in all use cases tested (Walter et al., 2019).

Ensemble methods combine predictions from multiple models to achieve superior performance compared to individual models alone. By leveraging synergies between these models, ensemble methods can enhance predictive performance and robustness. There are various methods for integrating these results, ranging from straightforward techniques such as union and intersection to majority voting, as well as more sophisticated approaches that involve optimization and machine learning models. Notably, ensemble methods can particularly benefit from incorporating multiple algorithms with diverse characteristics, as this diversity can lead to more accurate and robust predictions (Brown et al., 2005; Wang et al., 2020; Kuncheva and Rodríguez, 2014). In the field of bioinformatics, it is common practice to combine the results from various tools, such as peak callers in chromatin immunoprecipitation sequencing (ChIP-seq) data (Schweikert et al., 2012) and variant callers in DNA-seq data (Wang et al., 2020). For example, (Wang et al., 2020) demonstrated with “SomaticCombiner” that consensus approaches can outperform individual variant callers with respect to the identification of somatic variants (Wang et al., 2020), while another ensemble approach was shown to surpass individual classifiers in tuberculosis prediction (Osamor and Okezie, 2021). To our knowledge, there is no ensemble algorithm for the analysis of 4C-seq data. Given the availability of multiple 4C-seq interaction calling algorithms with distinct characteristics, we propose applying ensemble methods to 4C-seq data to improve interaction calling accuracy and robustness, ultimately leading to a more comprehensive understanding of the chromatin landscape and leveraging individual biases of the methods.

## Material and Methods

### Data acquisition

To identify 4C-seq datasets with validated interactions, we conducted a comprehensive literature research on PubMed, focusing on experiments from 2017 to 2025. We specifically targeted datasets that contained at least two conditions and two replicates each, originated from *Mus musculus* or *Homo sapiens*, and included at least one validated interaction. Interactions were considered validated if they were described by the authors as “significant interaction” and were biologically validated. Potential validation techniques are reciprocal 4C-seq experiments or other techniques such as CTCF-binding assays or ATAC-seq experiments (Kerschner et al., 2024; Hintermann et al., 2022). These validated interactions were used as the ground truth. The viewpoint and interaction positions were covered with a window of ±1,000 base pairs and exported to BED files. If in an experiment a reference genome older than mm10 or hg19 was used, we converted the genomic locations to mm10 or hg19 using UCSC liftOver (Hinrichs et al., 2006). To mitigate biases in cross-validation, we treated experimentally dependent datasets as groups to reduce the impact of experimental dependencies.

### Data processing

To develop an ensemble algorithm, training data were processed and interaction calling was performed with the available 4C-seq tools. To this end, we established a Snakemake pipeline (Köster and Rahmann, 2012) featuring quality control, preprocessing, interaction calling with peakC, r3Cseq, fourSig, and 4C-ker, as well as postprocessing steps. An overview of the development workflow is shown in Figure 1. Given the deprecated status of FourCSeq and in combination with the tools overall low recall in the benchmarking of Walter et al. (2019) and the current study, we excluded FourCSeq from the pipeline.

**Fig. 1.**
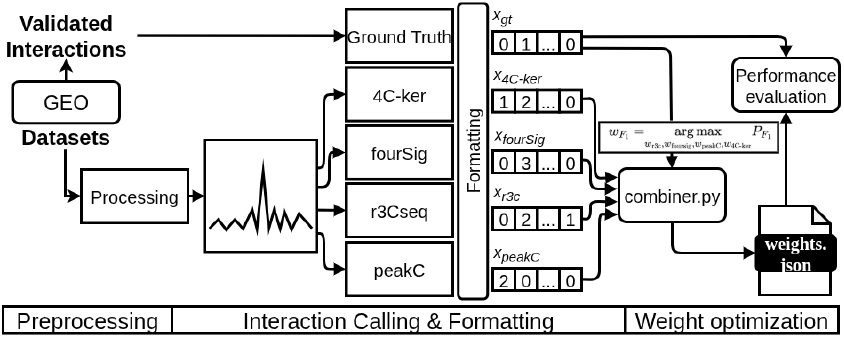
Schematic workflow of ensemble algorithm development exemplary for *F*_1_ score.

#### Preprocessing

During preprocessing, 4C-seq data is prepared for interaction calling and quality control is performed to ensure adequate data quality like sufficient fragment coverage. A detailed description of the preprocessing can be found in the supplementary 3.1.

#### Interaction calling

In this step, interaction calling is performed using r3Cseq, fourSig, peakC, and 4C-ker, which form the base of our ensemble algorithm and are referred to below as “base callers”. r3Cseq, peakC, and fourSig provide options to adjust the window size parameter. For r3Cseq, analyses were conducted using window sizes of 2k, 5k, and 100k base pairs. peakC was executed using window sizes of 11, 21, 31, and 51 fragments, while fourSig was run with window sizes of 1, 3, 5, and 11 fragments, respectively. These window sizes were chosen based on a previous benchmarking of 4C-seq algorithms (Walter et al., 2019).

#### Postprocessing

During postprocessing, the identified candidate interactions are collected and formatted to standardize the heterogeneous output formats. The formats differ in terms of the way they describe the interacting regions, the characterization of the significance of those, and the handling of replicates.

Since fragments are the smallest units used in a 4C-seq analysis, we map the results to the VFL. Due to the lack of validated cis and trans interactions, we limited our analysis to the near-bait area. Specifically, we defined the near-bait area as the region spanning ±1,500 fragments surrounding the viewpoint.

All base tools report the significance levels of interacting regions in different ways. fourSig and 4C-ker categorize peaks into distinct classes, r3Cseq outputs interacting regions detected in both experiment and control, quantified by q-values. peakC returns a vector of fragment ends which are considered to be peaks.

In order to develop an ensemble algorithm, we opt to harmonize the disparate significance information into discrete categories characterized by distinct scores. These scores categorize near-bait fragments into four classes: high-confidence candidate interactions (score: 3), candidate interactions (score: 2), low-confidence candidate interactions (score: 1), and non-interacting regions (score: 0). The results of the tools are mapped into those classes as described in the supplement 3.2.

Overall, the postprocessing includes fragment mapping, significance score homogenization, and replicate integration. The resulting data contains fragment start and end positions along with significance scores for both condition and control and is exported into BED files. Each row in these files represents an individual fragment with associated significance information, serving as input for downstream analyses.

### Optimal Weighted Calling

We combined four established 4C-seq interaction callers (r3Cseq, fourSig, peakC, and 4C-ker) to develop an ensemble interaction calling approach using a weighted voting strategy. To this end, we extracted nearbait significance scores (0, 1, 2, 3) from pipeline-generated BED files and represented them as score vectors *x*_r3c_, *x*_foursig_, *x*_peakC_, *x*_4C-ker_ for a given dataset *D*. Then, the ensemble score *S* is defined as

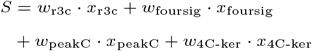

where the weights *w*_*i*_ *∈* [0, 0.5] are assigned to each base callers. Because *S* is dependent on the chosen dataset, its distribution varies across different data collections. To transform *S* into a discrete decision rule for interaction calling, we applied a decision threshold. For our analyses, this threshold was set at 0.5, ensuring that a significant result from even a single caller is sufficient to trigger a positive interaction call.

The validated interactions *x*_gt_ served as the ground truth for both weight optimization and performance benchmarking. Depending on the target metric (either *F*_1_ score or area under precision-recall curve (AUPRC)), the performance of the ensemble score *S* is defined as

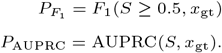

While AUPRC was calculated using the PROC package (pro), the *F*_1_ score was determined using the scikit-learn library (Pedregosa et al., 2011).

The performance metrics 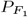 and *P*_AUPRC_ serve as objective functions for the weight optimization problem. To determine the optimal weights for each caller, we employed gradient-free optimization techniques and defined the optimization problem as:

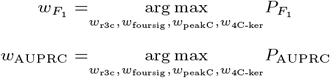

After benchmarking several algorithms within the Nevergrad library across varying iteration counts, the Covariance matrix adaptation evolution strategy (“CMA-ES”) and “OnePlusOne” (Rapin and Teytaud, 2018) showed the most reliable convergence for optimizing *F*_1_ and AUPRC, respectively. Consequently, these algorithms were selected to determine the final weights, using a budget of 100,000 trials per optimization loop (Fig. S2, more details about benchmarking: 3.4).

#### Global Weighting and Stability

As noted previously, the optimal weights 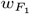 and *w*_*AUPRC*_, respectively, are inherently dependent on the underlying dataset *D* and may reflect specific dataset-caller biases. To derive a more robust and generalized weight, we sought a single global weight vector 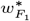 and 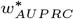, which remains effective across diverse data collections. By maximizing the averaged performance across *n* distinct datasets *D*_1_, …, *D*_*n*_, the optimization objective is reformulated as

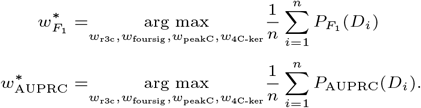

We compared the dataset-specific optimal weight with the global weights 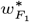and 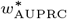 using a correlation analysis. To evaluate the stability and generalizability of the model to unseen data, we performed cross-validation using a leave-one-group-out (LOGO) routine to estimate generalization performance on unseen groups. Experimental dependencies, such as reciprocal viewpoints, were grouped to prevent information leakage during the validation process. Specifically, we quantified the impact of the LOGO procedure on both the overall performance as well as actual weight changes.

#### Ensemble Extension

Since the optimization is gradient-free, the objective extends readily to additional callers. Although the four base tools already include all widely used 4C-seq callers, a larger ensemble can still be built by running each tool multiple times with different hyper-parameter settings. Notably, window size is a critical parameter in interaction calling, as demonstrated by Walter et al. (2019), and significantly impacts tool performance. To account for this, we opted to represent each tool with multiple window size configurations within the optimized function. However, for clarity and conciseness, each tool is listed only once in the main text. The comprehensive set of functions, including all window size configurations, can be found in the supplementary materials (4).

### fourSynergy

We developed a framework called *fourSynergy*, to allow an easy application of our weighted voting based ensemble algorithm. *fourSynergy* consists of a Snakemake pipeline comparable to the pipeline we used to prepare the training data for the optimization, and an R Bioconductor package. In addition, it features an R Shiny application for easy and user-friendly access to the ensemble algorithm.

### Differential interaction analysis

The spatial organization of the genome can be altered in various diseases (Schuetzmann et al., 2018; Ghasemi et al., 2021). A key application of 4C-seq data analysis is the identification of spatial alterations between different conditions, such as a comparison of disease states with healthy controls. To this end, a differential interaction analysis can be performed.

Currently, only two 4C-seq algorithms, 4C-ker and FourCSeq, support differential analysis. Both tools use DESeq2, which was originally developed for differential RNA-seq analysis (Love et al., 2014). *fourSynergy*, also facilitates differential interaction analysis. Given that 4C-seq read counts follow a negative binomial distribution (Raviram et al., 2016), differential analysis is also conducted with DESeq2. The bases of this analysis are regions identified by our consensus interaction calling in condition and control. From these regions, the read counts are extracted and analyzed using DESeq2. Differential interactions are considered statistically significant if the adjusted p-value is *<* 0.05. To test this approach, we used our set of 20 datasets with validated differential interactions and evaluated the performance using the *F*_1_ score and AUPRC.

## Results

### Datasets

In total, our comprehensive literature review covered 108 4C-seq experiments and resulted in a set of 20 curated experiments, each containing at least one validated interaction. Of these, 9 datasets belong to experimentally related experiments and were therefore considered as groups in LOGO cross-validation (Tab. S1). These datasets serve as training data for developing the weighted voting-based ensemble algorithm, as well as ground truth to benchmark its performance against individual 4C-seq algorithms.

### Evaluation of 4C-seq analysis tools

The 4C-seq analysis tools r3c-seq, peakC, fourSig, and 4C-ker form the basis of the presented ensemble approach. Extensive benchmarking of these tools in various use cases on real and simulated 4C-seq data has already been performed (Walter et al., 2019). In order to compare the ensemble algorithm’s performance to those of the integrated base tools, we performed a benchmarking using 20 4C-seq experiments (Fig. 2). fourSig achieved mean *F*_1_ scores ranging from 0.02 to 0.04, which is comparable to 4C-ker (0.04) (see S2 for detailed statistics). In contrast, r3Cseq consistently achieved higher mean *F*_1_ scores, between 0.12 and 0.16, between the different window sizes that were tested. peakC yielded scores that highly depended on the window size: 0.18 (1 bp), 0.19 (21 bp), 0.12 (31 bp), and 0.06 (51 bp). These patterns were mirrored in the AUPRC. fourSig showed low AUPRC values (0.01–0.02), 4C-ker performed slightly better with an AUPRC of 0.02. r3Cseq attained higher AUPRC values, ranging from 0.07 to 0.08 across the tested window sizes, while peakC achieved the highest AUPRCs in the smaller window sizes (0.16 for 11 bp, 0.12 for 21 bp, 0.07 for 31 bp, and 0.03 for 51 bp). Overall, r3Cseq and peakC outperformed fourSig and 4C-ker in both *F*_1_ score and AUPRC, indicating that the interaction calling of this tools is more reliable in the tested area.

**Fig. 2.**
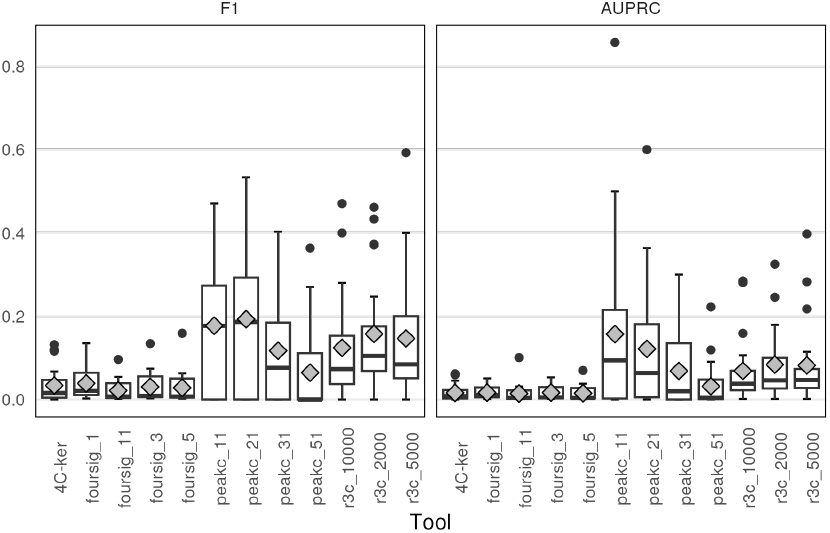
Predictive performance of individual 4C-seq algorithms. Boxplots illustrating the distribution of *F*_1_ scores and area under the precision-recall curve (AUPRC) values (y-axis) across different 4C-seq tools (x-axis), with grey diamonds indicating the mean values.

### Evaluation of single dataset-based solutions

To assess whether there is a more effective solution than using one of the base algorithms, we evaluated the performance of the best possible weighted voting per dataset according to optimization. Overall, the *F*_1_ and AUPRC optimized weighting solutions showed a higher predictive performance than the individual algorithms in both *F*_1_ and AUPRC (mean *F*_1_: 0.52, mean AUPRC: 0.38) (Fig. 3A). Notably, the optimal weightings vary depending on the performance metric used for optimization and exhibit distinct distributions across datasets (Fig. S3). Our findings suggest that weighted-voting solutions can achieve predictive performance higher than the base tools and raise questions about the existence of a global weighting that can be applied across multiple datasets.

**Fig. 3.**
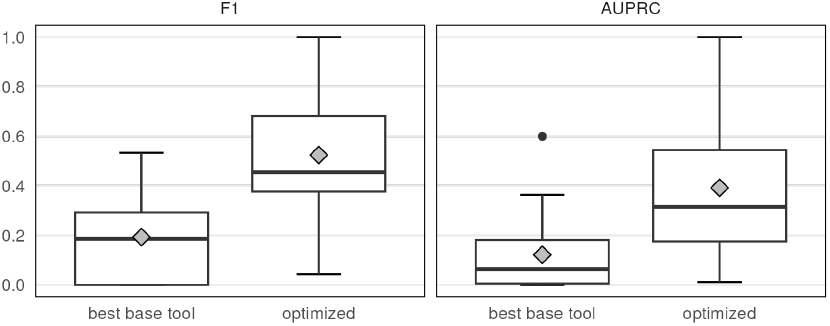
Predictive performance of indiviual 4C-seq algorithms and ensemble optimized on single datasets. Boxplots showing the distribution of *F*_1_ score and AUPRC values (y-axis) with grey diamonds indicating the mean values for the best performing single tool (peakC 21) and an ensemble solutions optimized on individual datasets (y-axis).

### Evaluation of multi dataset-based solution

Building on our previous results, we are trying to find out whether we can find a global solution for interaction calling by optimizing the tool weights in our consensus function across multiple datasets. The best-performing base tool (*F*_1_: peakC 21, AUPRC: peakC 11) achieved a mean *F*_1_ score of 0.19 and an AUPRC of 0.16. In contrast, the optimized solutions over all datasets yielded a mean *F*_1_ score of 0.34 and an AUPRC of 0.36 (Fig. 4A), outperforming the base tools in both evaluated predictive performance metrics. Furthermore, we observed that optimal weightings vary depending on the metric used for optimization (Fig. S4).

**Fig. 4.**
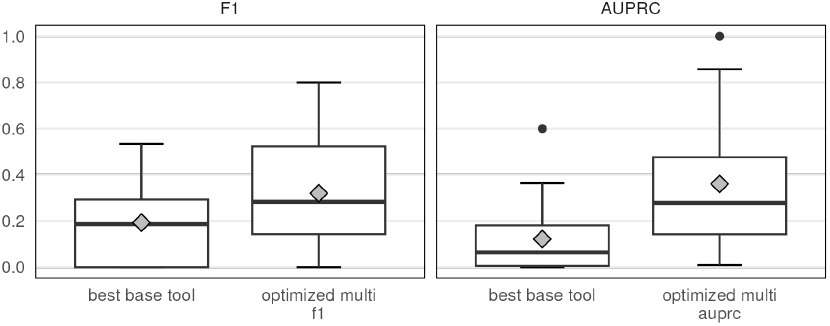
Predictive performance of individual 4C-seq algorithms and ensemble optimized on multi datasets. Boxplots showing the distribution of *F*_1_ score and AUPRC values (y-axis) for the best performing single tool (peakC 21) and an ensemble solutions optimized on all datasets (y-axis). Grey diamonds are representing the mean values.

### Cross validation

To evaluate the robustness of our multi-dataset solutions and to address the challenge of selecting the most suitable base tool for an unknown dataset, we employed LOGO. By applying LOGO to the ensemble solution and single tools, we compared the performance of the top-performing tool for each group on a held-out dataset. The results show that the ensemble approach outperforms the best single tools in LOGO, achieving a higher mean *F*_1_ score of 0.31 and AUPRC of 0.34, compared to 0.13 and 0.16, respectively (Fig. 5A). Depending on the predictive performance metric used for optimization, different tools were rated highest. For example, *F*_1_-optimized solutions for r3Cseq with a 2,000 bp window were highest weighted, while peakC solutions had the highest weights when optimized for AUPRC (Fig. S5).

**Fig. 5.**
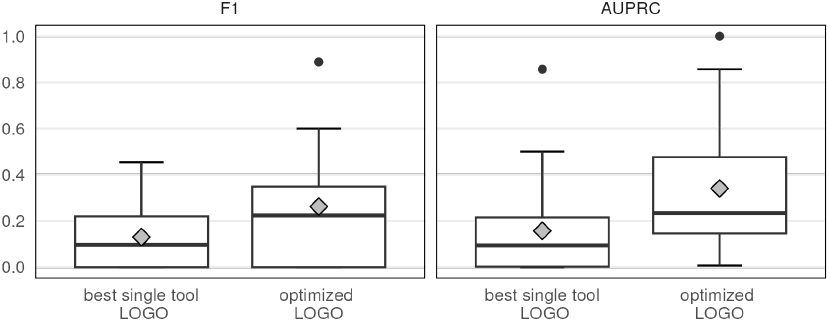
Predictive performance of single tool and ensemble solutions with Leave-One-Group-Out cross-validation (LOGO). A Boxplots showing the distribution of *F*_1_ scores and AUPRC values (y-axis) for the best-performing single tool with LOGO and ensemble solutions optimized on multiple datasets and validated with LOGO. The mean values are depicted by the gray diamonds.

### Differential analysis

The analysis of differential interactions is a useful application in 4C-seq data. Our results indicate that fourSynergy is able to identify differentially interacting regions and is able to outperform FourCSeq and 4C-ker in this task in the *F*_1_ score and AUPRC (Fig. S6).

### Runtime

r3C-seq was the fastest tool and fourSig the slowest. As expected, fourSynergy required more time than the individual tools (Fig. S7), with an additional pipeline overhead of ∼64 min on the tested dataset. Benchmark details are provided in subsection 3.3.

## Discussion

Our results demonstrate that ensemble algorithms outperform single algorithms in the task of interaction calling on 4C-seq data, highlighting the potential of ensemble algorithms in the analysis of next-generation sequencing data. In line with the broader idea of combining different tools during analysis (Schweikert et al., 2012; Wang et al., 2020), fourSynergy employs an approach using gradient-free optimization to determine optimal tool weights. This surpasses traditional majority voting methods, allowing fourSynergy to outperform individual tools in diverse scenarios. In particular, fourSynergy yields solutions that consistently outperform individual tools, even when evaluated through cross-validation, thereby providing a more reliable and accurate framework for interaction calling.

### Dataset structure

Another significant contribution of this study is the curation of 4C-seq datasets with validated interactions, which, to our knowledge, represents the largest collection compiled to date. This comprehensive dataset collection serves as a valuable resource for both training the ensemble algorithm and performance evaluation. During the literature review and the screening of 108 datasets, we identified a major challenge in the documentation of important experimental information, as well as insufficient data quality and problems with interaction validation. Despite these challenges, we were able to identify 20 curated datasets used to train the ensemble algorithm and evaluate its performance (Supplementary table S1). Although our dataset collection is extensive, it may not capture the full complexity and diversity of chromatin interaction patterns. Therefore, additional high-quality datasets could enhance the accuracy and robustness of our approach, leading to more reliable predictions and a better understanding of chromatin interactions.

Since biological validation of interactions is time- and cost intensive, usually only a few interactions can be validated and most of these datasets primarily comprise validated interactions, while lacking validated non-interactions. This could also be based on the fact that mostly only specific regions of interest, which are likely to harbor chromatin interactions, are validated or in the validation non-interacting regions might not always be reported in the publications. This scenario is reminiscent of PU learning scenarios (learning from positive and unlabeled data), in which only positive examples (validated interactions) are available, while negative examples (validated non-interactions) are rare or absent (Bekker and Davis, 2018). The absence of validated non-interactions has also consequences for our performance evaluation, as it restricts our ability to accurately estimate true negatives, making it difficult to distinguish between true negatives and false positives in the confusion matrix.

Another challenge in the training data is that a maximum of eight interactions were validated in a single experiment, with most datasets having only one validated interaction. This has implications for our confusion matrix, as there is a high chance that some real interactions were not validated, resulting in true positives being misclassified as false positives. This can lead to highly imbalanced ground truth data and incorrect annotations in the confusion matrix, which can affect our performance evaluation.

### Performance evaluation

The peculiarities of the data in terms of imbalance in ground truth data and potential misclassifications in the confusion matrix must be taken into account during performance evaluation. Recall should not be affected by these misclassifications, as it only represents the ratio of true positives to the sum of true positives and false negatives. Precision, on the other hand, can be affected by this, as the misclassification of true positives as false positives can lead to an underestimation of precision. To address this, *F*_1_ and AUPRC were used to evaluate the predictive performance of our algorithms. These metrics were used to provide a comprehensive assessment that reflects different aspects of the algorithm’s performance. While the *F*_1_ score offers a straightforward measure of predictive performance, it has some limitations in scenarios where precision and recall are not equally important. In contrast, the AUPRC provides a more nuanced view of an algorithm’s performance, capturing the trade-off between precision and recall across various thresholds. By combining these two metrics, we can gain a more complete understanding of our algorithms’ strengths and weaknesses, particularly in applications such as 4C-seq interaction calling.

### Combination strategy

There are various strategies to combine base algorithms in an ensemble approach. These strategies range from simple methods, such as intersection and union, to more sophisticated techniques, such as complex machine learning models. We performed analyses to assess the ensemble’s potential and the diversity among tools (see Supplementary 3.5)(Kuncheva, 2004). Overall, our results suggest that, although diversity among the top-ranked tools is limited, a simple aggregation of the five best-performing tools can yield performance gains (see Supplementary 3.5). In our case, we have a small number of imbalanced classes to classify. For such scenarios, Kuncheva and Rodríguez (2014) recommend weighted voting. Unlike a traditional majority vote, which assigns equal importance to all tools, weighted majority voting assigns higher weights to more reliable “expert” tools and can therefore further improve performance. Moreover, our analyses indicate that ensemble performance depends on the trade-off between complementarity and the predictive quality of the included tools (see Supplementary 3.5). Our weighted voting approach incorporates this trade-off directly into the optimization objective. In addition, the weighted voting approach can represent simpler operations, such as intersection, union, and majority votes, while also enabling more flexible and complex aggregation schemes.

### Consensus Approach for Cis and Trans Interactions

Our consensus approach is currently designed for near-bait analysis. The reason for this focus is the present lack of datasets with validated far-cis and trans interactions. Due to this, we were unable to optimize our consensus function in these areas. The transferability of our results to more distant cis or trans regions might be limited, as the behavior of the tools varies with distance from the viewpoint (Walter et al., 2019). Integration of information about the distance to the viewpoint and the implementation of distance-dependent weighting could improve the performance of the consensus approach and would make it more versatile. However, this would require datasets with validated interactions that feature diverse distances to the viewpoint.

### Runtime

Although the pipeline’s runtime is higher, its overhead is manageable and its benefits, including eliminating the need for time-consuming environment setup and providing comprehensive quality control, might outweigh the additional computational cost. Additionally, the small size of the 4C-seq datasets results in a comparatively short absolute runtime, mitigating the impact of increased computational costs.

### Outlook

This study demonstrates the potential of ensemble methods to surpass individual tools in the analysis of next-generation sequencing data. Our methodology, which uses a weighted voting approach based on gradient-free optimization, has the potential to be transferable to other NGS techniques. Furthermore, our approach could be extended to incorporate additional layers of complexity, such as spatially adapted weighting, to further enhance performance. For example, by adjusting the weights based on the distance to the viewpoint, our method could potentially capture more nuanced patterns in 4C-seq data. However, additional datasets that feature validated interactions with varying distances from the viewpoint would be required to fully exploit the potential of ensemble algorithms.

## Conclusion

Our study does not only provide a curated collection of 4C-seq datasets, but also demonstrates that weighted voting-based ensemble approaches outperform individual 4C-seq algorithms in predictive performance across a range of metrics. The optimal solutions vary depending on the metric used for optimization. To facilitate widespread adoption and ease of use, we have integrated our ensemble algorithm into a user-friendly framework called *fourSynergy*, which includes a Snakemake pipeline for reproducible and scalable analysis, as well as an R Bioconductor package and a Shiny app for intuitive and accessible data analysis.

## Supporting information

Supplementary fourSynergy

## Author contributions statement

Must include all authors, identified by initials, for example: G.H. and J.D. did the literature research. L.P., D.H. and S.M.W. worked on the weight optimization. S.M.W. implemented the Snakemake workflow. G.H. and S.M.W. implemented the R package. C.W., D.H., L.P. and S.M.W. wrote and reviewed the manuscript.

## Acknowledgments

The authors thank the anonymous reviewers for their valuable suggestions. This research was supported by the Open Access Publication Fund of the University of Muenster and “Innovative Medizinische Forschung” (IMF)(I-WA 11 23 15).

