## Supplementary fourSynergy for "fourSynergy: Ensemble-based interaction calling on 4C-seq data using gradient-free optimization"

#### 1 Supplementary Figures

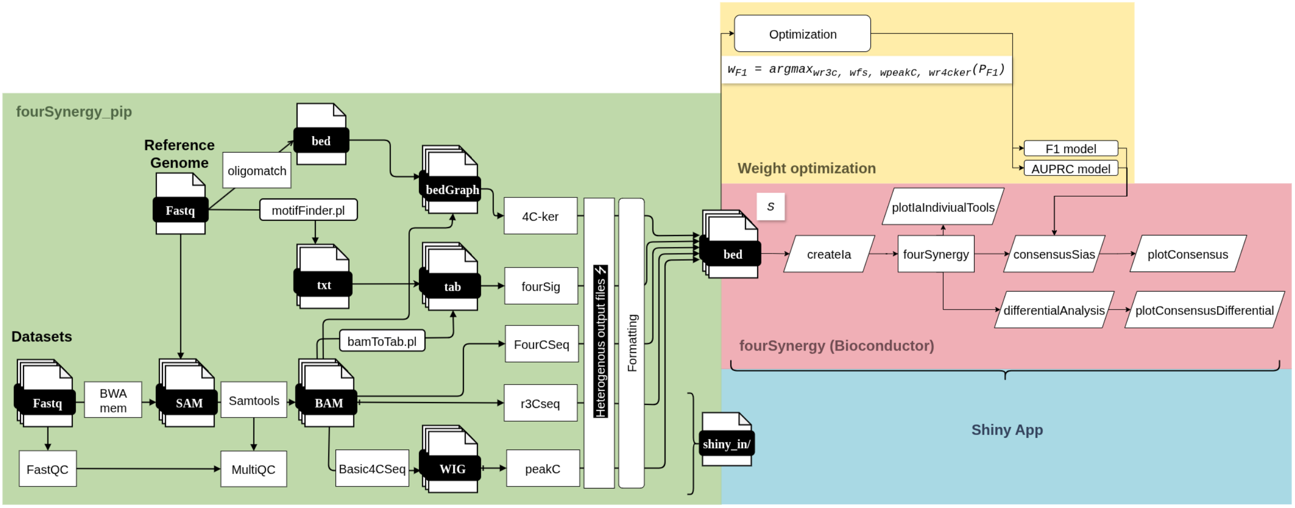


Figure S1: **Schematic overview of fourSynergy framework**. The framework features preprocessing pipeline, weight optimization, R Bioconductor package and R shiny application.

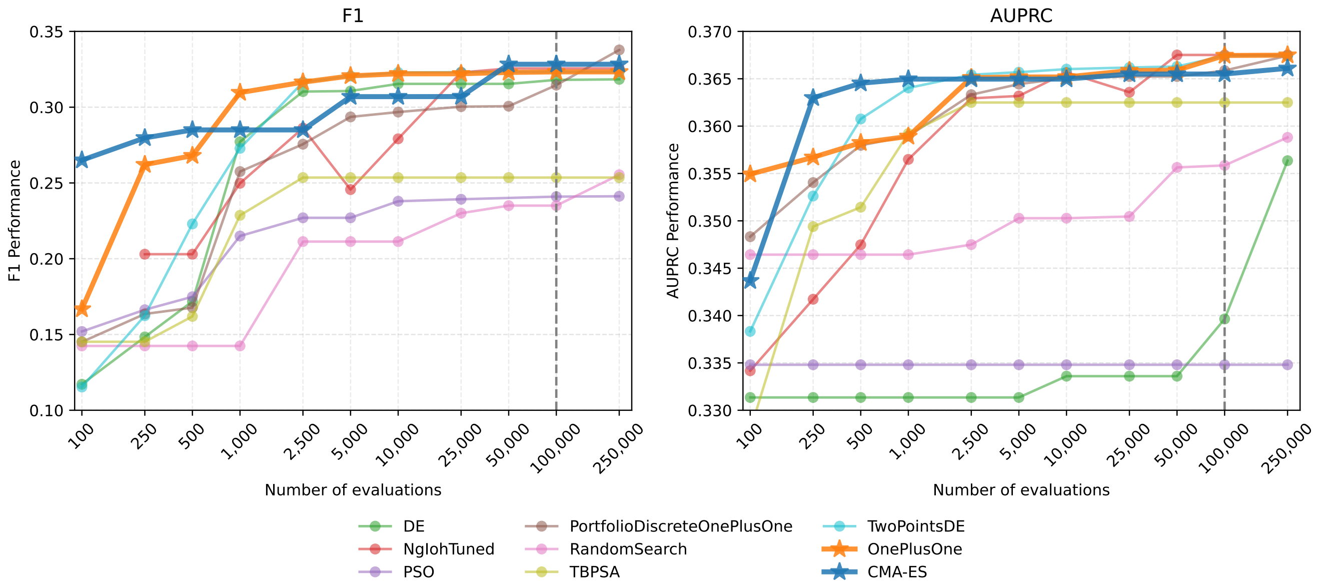


Figure S2: **Benchmarking Nevergrad optimizers.** Benchmarking Nevergrad optimizers across different iteration budgets using F1 score and area under precision recall curve (AUPRC).

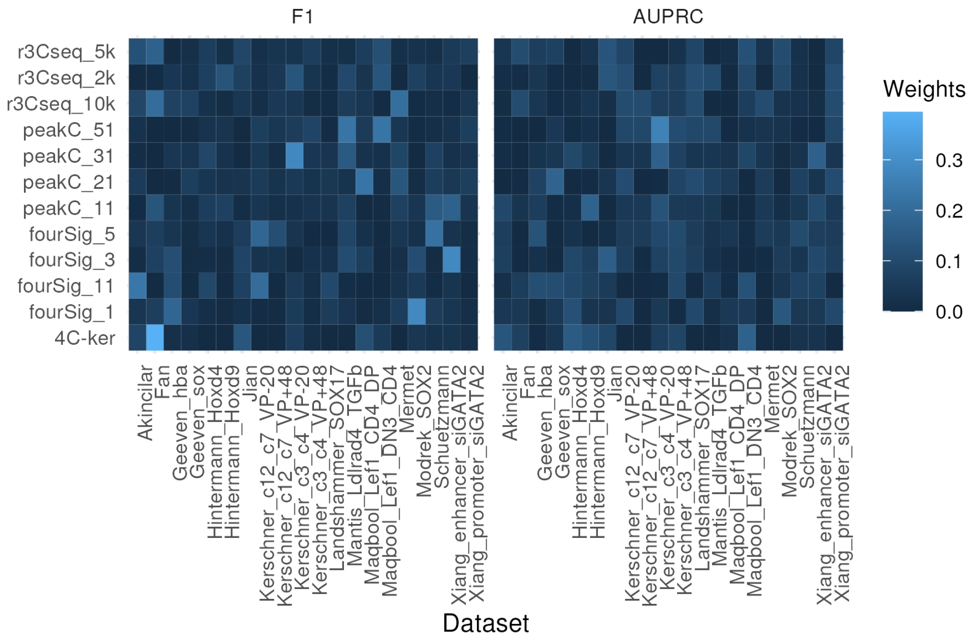


Figure S3: **Optimized weights for single dataset.** Heatmap showing the optimized weights for each tool (y-axis) across all datasets (x-axis), with color intensity indicating the weight.

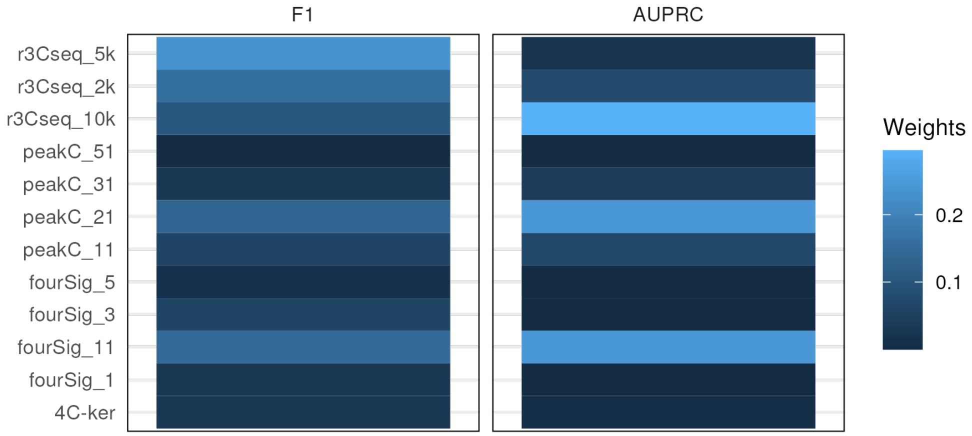


Figure S4: **Optimized weights over all datasets.** Heatmap showing the optimized weights for each tool (y-axis) over all datasets (x-axis), with color intensity indicating the weight.

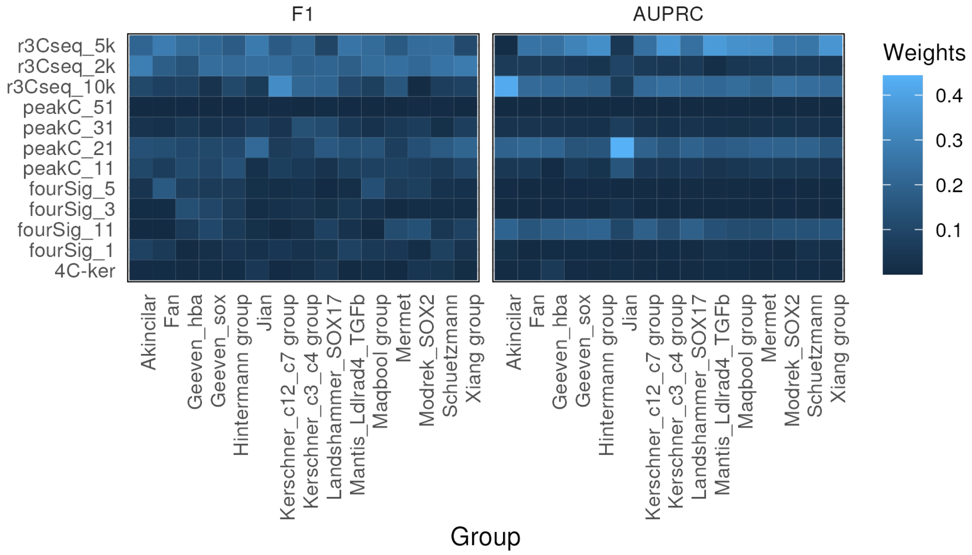


Figure S5: **Optimized weights for Leave-One-Group-Out cross validation.** Heatmap showing the optimized weights for each tool (y-axis) across all datasets groups (x-axis), with color intensity indicating the weight.

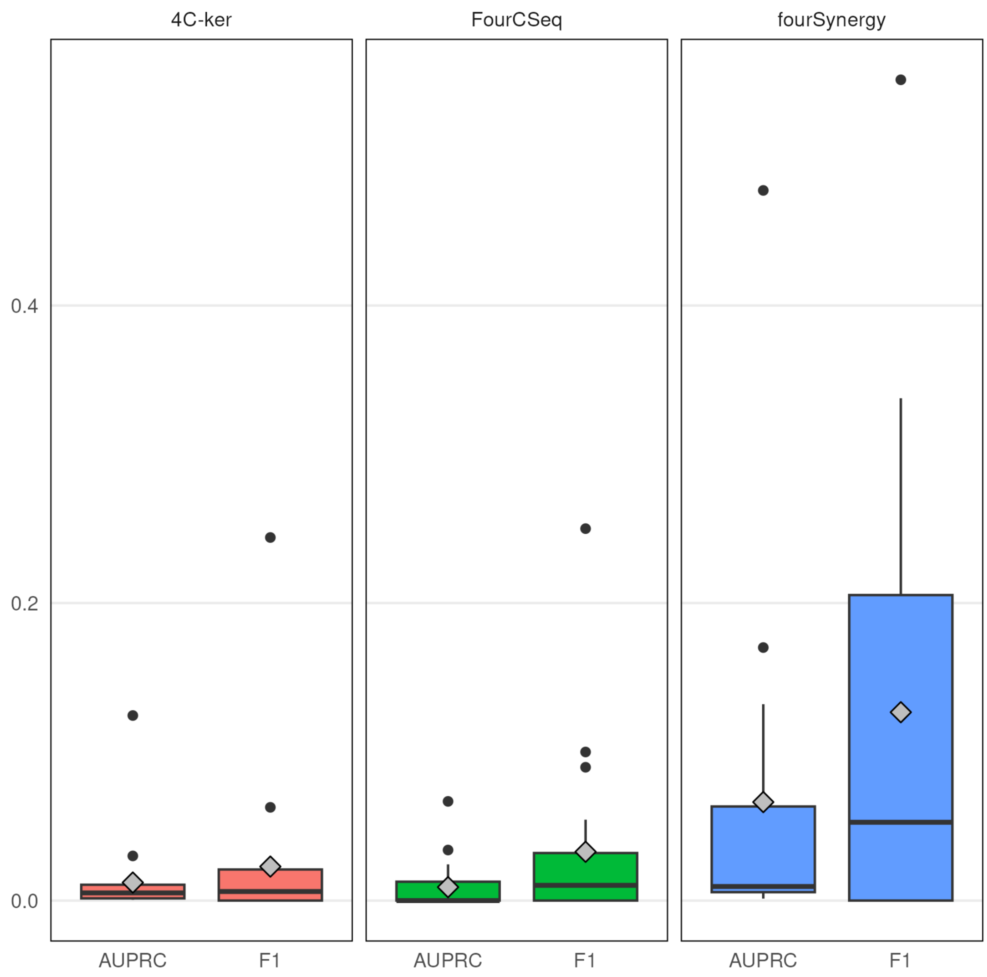


Figure S6: **Predictive performance of differential interaction calling using different 4C-seq algorithms.**

Boxplots illustrating the distribution of F1 scores and area under the precision-recall curve (AUPRC) values (y-axis) across different 4C-seq tools (x-axis), with diamonds indicating the median values.

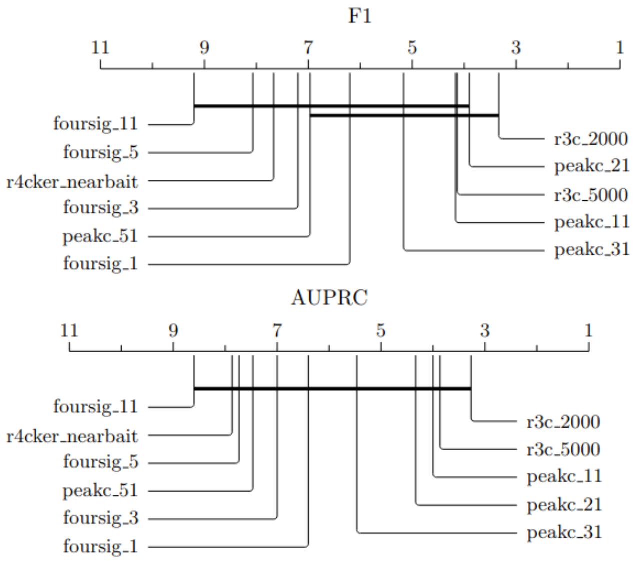


Figure S8: **Critical difference analysis of base tools.** Critical difference analysis of the predictive performance of the base tools across the curated collection of 4C-seq datasets (n=20n=20n=20).

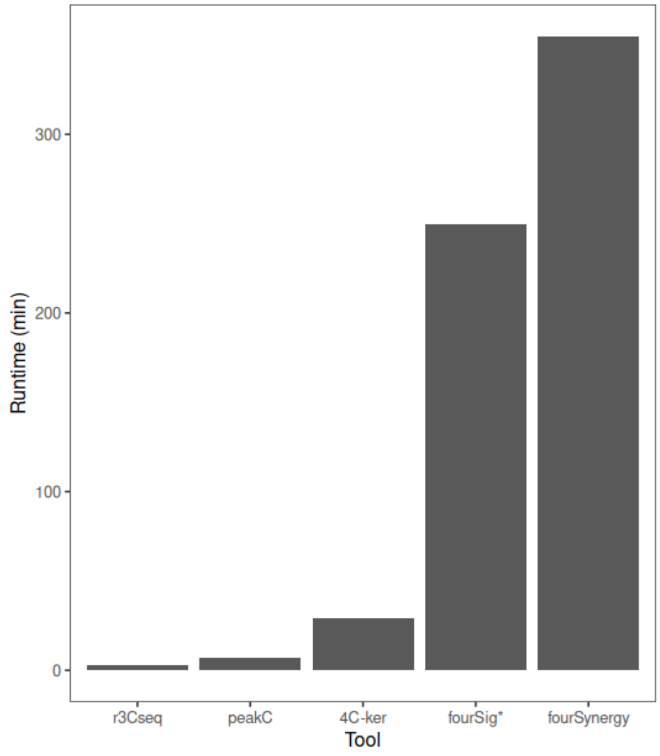


Figure S7: **Runtime Benchmarking of 4C-seq analysis tools**. Runtime of base tools (x-axis) in minutes (y-axis). Runtime includes preprocessing and tool execution with window sizes mentioned in Materials and Methods.

#### 2 Supplementary Formulas

$$S = w_{\text{r3c 2k}} \cdot x_{\text{r3c 2k}} + w_{\text{r3c 5k}} \cdot x_{\text{r3c 5k}} + w_{\text{r3c 1k0}} \cdot x_{\text{r3c 1k0}} + \tag{1}$$

$$w_{\text{foursig 1}} \cdot x_{\text{foursig 1}} + w_{\text{foursig 3}} \cdot x_{\text{foursig 3}} + w_{\text{foursig 5}} \cdot x_{\text{foursig 5}} + w_{\text{foursig 11}} \cdot x_{\text{foursig 11}} + \tag{2}$$

$$w_{\text{peakC 11}} \cdot x_{\text{peakC 11}} + w_{\text{peakC 21}} \cdot x_{\text{peakC 21}} + w_{\text{peakC 31}} \cdot x_{\text{peakC 31}} + w_{\text{peakC 51}} \cdot x_{\text{peakC 51}} + \tag{3}$$

$$w_{\text{4C-ker}} \cdot x_{\text{4C-ker}} \tag{4}$$

where

$$\begin{aligned} w_{\text{r3c 2k}}, w_{\text{r3c 5k}}, w_{\text{r3c 10k}}, \\ w_{\text{foursig 1}}, w_{\text{foursig 3}}, w_{\text{foursig 5}}, w_{\text{foursig 11}}, \\ w_{\text{peakC 11}}, w_{\text{peakC 21}}, w_{\text{peakC 31}}, w_{\text{peakC 51}}, \\ w_{\text{4C-ker}} \in [0, 0.5] \end{aligned}$$

$$w^*_{F_1} = \underset{(w_{\text{r3c 2k}}, w_{\text{r3c 5k}}, w_{\text{r3c 10k}}, w_{\text{foursig 1}}, w_{\text{foursig 3}}, w_{\text{foursig 5}}, w_{\text{foursig 11}}, w_{\text{peakC 11}}, w_{\text{peakC 21}}, w_{\text{peakC 31}}, w_{\text{peakC 51}}, w_{\text{4C-ker}})}{\arg\max} \left( \frac{1}{n} \sum_{D_0, \ldots, D_n} P_{\text{F1}} \right)$$

### 3 Supplementary Methods

#### 3.1 Preprocessing

The entry point of the pipeline are FASTQ files downloaded via the Gene Expression Omnibus (GEO) [1]. Relevant information on the 4C-seq experiment is provided in a config file.
In the first step of preprocessing, initial quality control of the FASTQ files is performed using FASTQC [2]. The reads are aligned to the reference genome using BWA MEM (0.7.18-r1243-dirty) [3]. The resulting BAM files are subsequently indexed, sorted, and the alignment statistics are summarized using Samtools [4]. All results are then collected by MultiQC [5]. Basic4CSeq further performs a 4C-seq-specific quality control and basic analysis [6]. In this step, the reference genome is fragmented at the restriction sites to create a virtual fragment library (VFL), in silico mimicking the restriction enzyme digestion performed in the wet lab. This is particularly important as the virtual fragment library forms the basis of our analysis. In addition, we perform various steps to convert the input data into the appropriate formats for each 4C-seq tool. The different interaction callers have varying input requirements. r3Cseq and 4C-ker operate on BAM files, while 4C-ker additionally needs a reduced genome file to map reads on. peakC utilizes WIG files, while fourSig requires TAB files, which can be created using scripts from the fourSig software suite. Detailed information regarding preprocessing can be found in the workflow in the supplementary materials (Fig.S1).

#### 3.2 Significance score mapping

As fourSig employs a similar categorical classification, its results could be seamlessly integrated into our significance scoring system [7]. For r3Cseq, on the other hand, we used the q-values to assign the fragments to our significance categories. Fragments with a q-value <0.01 are considered to be high-confidence candidate interactions (3), those with a q-value >= 0.01 and <0.05 to be candidate interactions (2), those with q-value >= 0.05 and <0.1 to be low-confidence candidate interactions (score: 1) and >= 0.1 to be non-interacting regions (0) [8]. peakC returns a vector of peak ends based on the given FDR (False Discovery Rate) parameter. To integrate these results into our significance scoring system, peakC is executed several times with varying FDR settings (0.01, 0.05, 0.1) [9]. Fragments are then classified into our significance scoring system using cutoffs analogous to those applied to r3Cseq. 4C-ker categorizes regions into three classes: ”high interactions”, ”low interactions”, and ”no interactions”. These classes are mapped to our significance scoring system as follows: ”high interactions” are assigned a score of 3, ”low interactions” a score of 2, and ”no interactions” a score of 0 [10].
peakC, r3Cseq and 4C-ker summarize the replicates in their outputs in analyses with multiple conditions and replicates. In contrast, fourSig does not offer integration routines for replicates. To ensure consistency across the tools outputs, we aggregated the interaction scores generated by fourSig across all replicates. An interaction was considered valid if it was detected in at least half of the replicates. If one of the tools fails or is not calling interactions, a dummy file is written and the significance scores are set to 0.

#### 3.3 Runtime benchmarking

The runtime was tested on a Linux system with an Intel(R) Xeon(R) CPU E5-2695 v4 @ 2.10GHz, running Debian 6.1.153-1, with 504 GB of total RAM, our pipeline utilized a maximum of 16 GB RAM during execution.

#### 3.4 Optimizer benchmarking

For selecting the most suitable optimizer, we benckmarked several Nevergrad implementations across increasing evaluation budgets to assess their effectiveness for the average multi-optimization task. Therefore, this benchmark does not constitute the cross-validation procedure, as performance is evaluated directly on the entire set. For each optimizer, performance was tracked as a function of the number of objective-function evaluations. The tests are not performed only once per metric and optimizer, but restarted for every budget step. Results were summarized using F1 score and AUPRC. This analysis was intended to compare how quickly different optimizers reached strong solutions and how performance evolved with additional iterations. Together, these benchmarks provide an overview of both convergence behavior and final optimizer choice for both metrics.

Figure S2 summarizes the results from the described analysis. Comparing F1 score and AURPC suggests that optimization with respect to F1 score is the more challenging problem, as most optimizers start below 0.2 and improve gradually over time. In contrast, AUPRC starts above 0.3 and reaches a similar maximum level at the end of the optimization. In addition, AUPRC remains relatively stable from an budget of 2500 onward. The analysis does not reveal a clear convergence point for either metric, as some optimizers still improves between 100,000 and 250,000 iterations. Nevertheless, for most of the best-performing optimizers, the improvement in this range is only marginal, which led us to select 100,000 iterations as a practical sweet spot for all subsequent analysis. Based on this benchmark, we choose the CMA-ES for all F1 score-related optimizations and OnePlusOne for the AUPRC-related optimizations, as these optimizers achieved the best performance at the budget of 100,000 iterations.

##### 3.5 Ensemble potential and diversity

We calculated theoretical upper bounds on ensemble performance following [11] for an ensemble comprising the five top-ranked tools identified from a critical-difference analysis (Fig. S8). Let $\{M_1, ..., M_5\}$ be our five models and let $k = (L+1)/2$ with $L$ the number of models (i.e., 5), then the upper bound is given by

$$maxP_{maj} = min\{1, \sum(k), \sum(k-1), ..., \sum(1)\} \tag{5}$$

with $\sum m = 1/m \sum_{i=1}^{L-k+m} p_i$ with $m = 1, ..., k$.

These tools were selected because they ranked consistency highly with respect to both $F_1$ and AUPRC. We compared this upper bound with the performance of the best individual tool according to the same critical-difference analysis (r3Cseq_2000). In 13 datasets, the upper bound of the ensemble exceeded the observed $F_1$ score of r3Cseq_2000. Averaged across datasets, the best-performing single tool (r3cseq_2000) achieved a mean F1 score of 0.16, whereas the corresponding theoretical upper bound for the top-five ensemble was 0.25. Furthermore, we quantified the diversity among the top five tools by measuring entropy [11]. A fragment was classified as a "call" if the significance score was >= 1.

$$entropy = \frac{1}{N}\frac{2}{L-1}\sum_{j=1}^{N} min\{(\sum_{i=1}^{L} y_{ij}), (L - \sum_{i=1}^{L} y_{ij})\} \tag{6}$$

The mean entropy was 0.049, indicating limited disagreement among the top-ranked tools.

Overall, these findings suggest that, despite relatively low diversity, the top-ranked tools still contain complementary information that could be exploited using ensemble-based methods. In particular, the gap between the best individual tool and the theoretical ensemble upper bound indicates that performance gains are already achievable with simple aggregation strategies such as majority voting. More flexible approaches, such as our weighted voting procedure, further improve performance by assigning greater influence to consistently stronger tools. Further analyzes demonstrated that increasing the number of integrated tools led to higher diversity but simultaneously to a lower theoretical ensemble upper bound. This suggests that ensemble performance depends on the balance between complementarity and the predictive quality of the included tools. In our weighted voting approach, this balance is incorporated directly into the optimization objective, allowing the method to determine both which tools should contribute to the final prediction and how strongly.

### 4 Supplementary Tables

| Dataset | organism | VP chromosome | conditionRep | controlRep | RE1 | RE2 | GEO | No. validated interactions | Group | Reference |
|---|---|---|---|---|---|---|---|---|---|---|
| Akincilar | human | 5 | 2 | 2 | HindIII | DpnII | GSE77265 | 5 | - | [12] |
| Fan | mouse | 5 | 2 | 2 | HindIII | DpnII | GSE87299 | 2 | - | [13] |
| Schuetzmann | mouse | 2 | 2 | 2 | NlaIII | NlaIII | GSE110683 | 2 | - | [14] |
| Geeven_hba | mouse | 11 | 3 | 3 | DpnII | Csp6I | GSE105177 | 2 | - | [9] |
| Geeven_sox | mouse | 3 | 3 | 3 | DpnII | Csp6I | GSE105177 | 1 | - | [9] |
| Mermet | mouse | 10 | 3 | 3 | DpnII | NlaIII | GSE101423 | 1 | - | [15] |
| Maqbool_Lef1_DN3_CD4 | mouse | 3 | 2 | 2 | DpnII | Csp6I | GSE144586 | 6 | Maqbool group | [16] |
| Maqbool_Lef1_CD4_DP | mouse | 3 | 2 | 2 | DpnII | Csp6I | GSE144586 | 4 | Maqbool group | [16] |
| Jian | human | 11 | 3 | 3 | BglII | NlaIII | GSE111863 | 1 | - | [17] |
| Modrek_SOX2 | human | 3 | 2 | 2 | DpnII | Csp6I | GSE94962 | 7 | - | [18] |
| Mantis_Ldlrad4_TGFb | mouse | 18 | 2 | 2 | NlaIII | DpnII | GSE197013 | 8 | - | [19] |
| Kerschner_c12_c7_VP-20 | human | 7 | 2 | 2 | NlaIII | DpnII | GSE220855 | 1 | Kerschner_c12_c7 group | [20] |
| Kerschner_c12_c7_VP+48 | human | 7 | 2 | 2 | NlaIII | DpnII | GSE220855 | 1 | Kerschner_c12_c7 group | [20] |
| Kerschner_c3_c4_VP-20 | human | 7 | 2 | 2 | NlaIII | DpnII | GSE220855 | 1 | Kerschner_c3_c4 group | [20] |
| Kerschner_c3_c4_VP+48 | human | 7 | 2 | 2 | NlaIII | DpnII | GSE220855 | 1 | Kerschner_c3_c4 group | [20] |
| Hintermann_Hoxd1 | mouse | 2 | 1 | 2 | DpnII | NlaIII | GSE195591 | 2 | Hintermann group | [21] |
| Hintermann_Hoxd4 | mouse | 2 | 2 | 2 | DpnII | NlaIII | GSE195591 | 2 | Hintermann group | [21] |
| Landshammer_SOX17 | human | 8 | 3 | 3 | NlaIII | DpnII | GSE178990 | 1 | - | [22] |
| Xiang_enhancer_siGATA2 | human | X | 2 | 2 | DpnII | NlaIII | GSE275926 | 1 | Xiang group | [23] |
| Xiang_promoter_siGATA2 | human | X | 2 | 2 | DpnII | NlaIII | GSE275926 | 1 | Xiang group | [23] |

Table S1: Overview of datasets and validated interactions

| Tool | Mean F1 | Median F1 | SD_F1 | Mean AUPRC | Median AUPRC | SD AUPRC |
|---|---|---|---|---|---|---|
| foursig_1 | 0.04 | 0.02 | 0.04 | 0.02 | 0.01 | 0.01 |
| foursig_11 | 0.02 | 0.01 | 0.03 | 0.01 | 0.00 | 0.02 |
| foursig_3 | 0.03 | 0.01 | 0.04 | 0.02 | 0.01 | 0.02 |
| foursig_5 | 0.03 | 0.01 | 0.04 | 0.02 | 0.00 | 0.02 |
| opo_indiv_auprc | 0.06 | 0.02 | 0.09 | 0.40 | 0.32 | 0.29 |
| opo_indiv_f1 | 0.52 | 0.45 | 0.26 | 0.38 | 0.31 | 0.29 |
| opo_multi_auprc | 0.12 | 0.07 | 0.13 | 0.36 | 0.28 | 0.30 |
| opo_multi_auprc_logo | 0.07 | 0.03 | 0.09 | 0.34 | 0.23 | 0.31 |
| opo_multi_f1 | 0.34 | 0.26 | 0.30 | 0.34 | 0.28 | 0.29 |
| opo_multi_f1_logo | 0.31 | 0.25 | 0.27 | 0.32 | 0.27 | 0.29 |
| peakc_11 | 0.18 | 0.18 | 0.16 | 0.16 | 0.09 | 0.21 |
| peakc_21 | 0.19 | 0.19 | 0.17 | 0.12 | 0.06 | 0.15 |
| peakc_31 | 0.12 | 0.08 | 0.14 | 0.07 | 0.02 | 0.09 |
| peakc_51 | 0.06 | 0.00 | 0.11 | 0.03 | 0.00 | 0.06 |
| r3c_10000 | 0.12 | 0.07 | 0.13 | 0.07 | 0.04 | 0.08 |
| r3c_2000 | 0.16 | 0.10 | 0.14 | 0.08 | 0.05 | 0.08 |
| r3c_5000 | 0.15 | 0.08 | 0.15 | 0.08 | 0.05 | 0.10 |
| r4cker_nearbait | 0.03 | 0.02 | 0.04 | 0.02 | 0.01 | 0.02 |
| single_tool_logo_auprc | 0.18 | 0.18 | 0.16 | 0.16 | 0.09 | 0.21 |
| single_tool_logo_f1 | 0.13 | 0.10 | 0.15 | 0.12 | 0.03 | 0.22 |

Table S2: Predictive performance statistics

[2] FastQC, Jun 2015.

[11] Ludmila I Kuncheva. *Combining Pattern Classifiers*. John Wiley & Sons, Ltd, 2004.
